# Awareness of MERS-CoV among Staff Members of Prince Sultan Military Medical City in Riyadh, Saudi Arabia

**DOI:** 10.1101/474205

**Authors:** Saud H Aman, Mohammed I Aljaber, Abdullah I Alwehaibi, Fahad H Aman, Hamad A Algaeed, Sultan M Almasoud, Majed A Alahmari, Omair Al Hussain, Ahmed Elhag

**Author notes:** Corresponding author: (SHA). This is the Senior Author (Saud H Aman). These authors contributed equally to this work (Hamad A Algaeed, Sultan M Almasoud, Majed A Alahmari). These authors also contributed equally to this work (Mohammed I Aljaber, Abdullah I Alwehaibi, Fahad H Aman, Omair Al Hussain, Ahmed Elhag).

## Abstract

**Background:** The Middle East respiratory syndrome coronavirus (MERS-CoV) developed infections that caused serious epidemics. Special priority of awareness and prompt initiative involvement of health workers (HW) during such intensified health situation necessitated an assessment of their preparedness, appropriate attitudes and protective strategies for better efficiency modes and acceleration during emergency.

**Methods:** HW of Prince Sultan Military Medical city in Riyadh, Saudi Arabia were reviewed through specifically designed questionnaires to acquire the demanded data. It included clinical and demographic information about the viral diseases, associated signs and symptoms, transmission and protection, and attitudes about the MERS-CoV disease.

**Results:** The study was accomplished between August and November, 2017 and 477 participants of the medical city, successfully completed the study questionnaire (Appendix I). Females represented a majority and there was an indirectly proportional decrease with the increasing age. Gradual educational increase levels reflected dominance of the university degree holders. Jobs were dominated by nurses and non-Saudis were a majority whilst, the highly experienced, (>10 years) were a minority. A majority recognized the viral transmission methods, popular information sources of MERS-CoV and associated medical terms. Highest scores were observed in dealing with protective aids and recognizing symptoms of disease. High adherence to hand hygiene protocols and correct washing steps were recorded. Correct and high levels were observed in taking preventive measures and avoiding infection. Participants responded correctly to negative and wrong actions that patients should refrain from. High scores were observed in taking appreciable attitudes towards oneself and towards others.

**Conclusions:** Expatriates were majority and nurses were dominant which, necessitates Saudization of this sector. Ministry of health pamphlets and seminars were of less impact in invigilating HW, hence, more attention and efforts are demanded. HW were quite aware of the basic and emergent health policies during epidemic episodes of MERS-CoV.

## Introduction

The coronavirus that causes acute respiratory infection in humans is known as the Middle East Respiratory Syndrome (MERS-CoV) and was discovered in 2012 [1].

Reports indicate that two-thirds of the infection that effect humans are not related to camels. Kharma and colleagues thus suggested that another species could act as an intermediary to human infection. Since fragments of the human MERS-CoV RNA sequence was isolated in a Taphozous bat [2], [3] they have since been suspected as a possible source of viral transmission to humans. However, the rare contact between humans and bats implies hypothetically that they are the least responsible for the human infection. For deeper investigation to find the role that bats play as MERS-CoV risk factor of viral transmission, more research is needed [4].

Human-to-human transmission is an increasingly public health concern since more than fifty percent of the confirmed secondary cases have been identified within hospitals and healthcare center boundaries [2]. Most of this coronavirus infections reported in Saudi Arabia were transmitted through human interactions within the health center boundaries, that might have been due to inadequate infection control within these areas [5].

Usually the incubation period of MERS-CoV after infection is within two weeks [6]. According to recommendations of the WHO and CDC, individuals who had visited the Middle East region within the previous 14 days should be screened for MERS-CoV [7]. In a hospital outbreak in September 2012, the first cases of pneumonia that were caused by the new coronavirus (MERS-CoV) were reported by the World Health Organization [8]. The reported symptoms in patients included fever, cough, shortness of breath and gastrointestinal symptoms, associated with a mortality rate of 65% [6]. Most patients infected with MERS-CoV show signs of pneumonia and ARDS, as well as acute kidney injury in some of the cases [9]. Whether the severity of MERS-CoV in patients depends on other conditions is still not clear [10]. One study in Saudi Arabia reported that an overwhelming majority of patients with MERS-CoV had underlying comorbidities [6]. Previous studies suggest that patients with diabetes, chronic lung diseases, renal failure, or immunocompromised are prone to greater risk of severe coronavirus onsets, however, there is little evidence to suggest that such underlying comorbidities contributed to disproportionate infection of MERS-CoV [8]. The WHO estimated the mortality rate of the coronavirus to be approximately 30% [2], [11]. Currently, the treatment is rather non-specific and there are no recommended antiviral agents [2]. Though, Nitazoxanide which is a broad-spectrum antiviral drug is being tested against influenza and other viral respiratory infections and it exhibited appreciable activity against MERS-CoV and other viruses by inhibiting their N protein expression [12], [13]. Since human-to-human transmission is said to occur in more than 60% of the total infections, mostly involving medical staffs, inpatients prone to and are at an increased risk of MERS-CoV infection. The WHO suggested that infection in healthcare settings were due to overcrowding and insufficient control measures [14]. Secondly, it would be necessary for the patients to avoid contact with camels or to eat properly cooked camel meat [15]. Sufficient knowledge of the etiology and prevention of the Middle East respiratory syndrome coronavirus among healthcare professionals is essential in combating the virus.

Significant improvement in patients administered with a combination of ribavirin and interferon alfa-2a was seen, in comparison with controls that received only supportive medical care [16]. However, further evaluation of the therapy regimen is required to enable the conclusive use of the therapy in the management of the disease.

Several preventive measures to control infections in healthcare settings, to expedite management of the disease is essential, such as developing of an emergency department visual triage scores for early detection of the MERS-CoV [17]. It has been shown that the use of triage for early detection of various types of disease had positive outcomes. Earlier studies had speculated this and suggested that it might play major roles in the early detection of this disease [18], [19].

Droplets from infectious sources are responsible for the infection, hence WHO recommendations are to follow standard precautions, when caring for patients suffering from acute respiratory problems [5]. These prevention measures that include protection of the nose, eyes, and any other places that the medical staff can contract the disease from when caring for known or suspected cases of the Middle East coronavirus infection, should be observed [20], [21]. CDC [22] recommends that those infected individuals who do not have inpatient support can be cared for in isolation in their homes.

People who are patients with diabetes, lung diseases, renal failure, or immunocompromised should avoid contact with camels or areas where camels are most likely to be found [5], [8]. In addition, the WHO recommends such patients to avoid undercooked camel meat, raw milk, and urine in addition to practicing proper hygiene.

Finally, screening for coronavirus at border points as well as the issuing travel advisories and human/ camel movement restrictions are recommended to avoid cross-border spreading and infection. Also, high levels of vigilance should be implied to prevent infection from persons returning from the middle east region [2].

This study aims at exploring the level of awareness about MERS-CoV among the Staff at Prince Sultan Military Medical City in Saudi Arabia. The knowledge generated from this study is expected to help staff improve their MERS-CoV awareness levels to enhance the fight against the problem.

## Materials and Methods

### Research Design

This study is a cross-sectional Study that used a survey to assess the awareness level of MERS-CoV among staff members of Prince Sultan Military Medical city in Riyadh, Saudi Arabia. A structured questionnaire on the disease was prepared in both English and Arabic, based on a review of the literature and consultation with regional community health teams in Saudi Arabia.

### Methods

The questionnaire was administered through the various departments in the health city and was distributed through manual approaches. The participants of this study were selected from the whole staff of the medical city, including all groups of staff workers. All participants in this manuscript has given written informed consent, approved by the IRB (Institutional Research Board and Research Ethics Committee) to publish this research study.A total of 570 responses were targeted to increase the reliability and validity of this study.

### Statistical analysis

The data collected from the participants were entered into SPSS software (version 20) for analysis. A range of descriptive and inferential statistics were carried out to explore the awareness of the staff about MERS-CoV.

## Results

There were 477 responders (with a response rate of 82.7%) of all participants who successfully completed the required information data required in this study questionnaire (Appendix I). These data were collected and compiled in tables. Table 1 reflected the demographic profile (categories, parameters, frequencies and percentages) of those participants, selected from staff members of Prince Sultan Military Medical city in Riyadh. Age of the participants was categorized in five range groups. The group range with the least count was <20 years (6, 1.3%). Next age range groups got an indirectly proportional decrease in count with increasing age, 20 – 29 years (191, 40.0%); 30 – 39 years (170, 35.6%); 40 – 49 years (78, 16.4%) and >50 years (27, 5.7%). There were 5 missing (1.0%) records. The majority (276, 57.9%) of these participants were females. With respect to nationality, Non-Saudis were a majority (349, 73.2%).

**Table 1.** Demographic profile (n=477)

| **Table 1. Demographic profile (n=477)** |  |  |  |
| --- | --- | --- | --- |
| **Category** | **Parameter** | **Frequency** | **%** |
| Age | <20 | 6 | 1.3% |
|  | 20 – 29 | 191 | 40.0% |
|  | 30 – 39 | 170 | 35.6% |
|  | 40 – 49 | 78 | 16.4% |
|  | >50 | 27 | 5.7% |
|  | Missing | 5 | 1.0% |
| Gender | Male | 183 | 38.4% |
|  | Female | 276 | 57.9% |
|  | Missing | 18 | 3.8% |
| Nationality | Saudi | 128 | 26.8% |
|  | Non-Saudi | 349 | 73.2% |
| Job | Non-health worker | 8 | 1.7% |
|  | Nurse | 240 | 50.3% |
|  | Specialist | 15 | 3.1% |
|  | Physician | 156 | 32.7% |
|  | Pharmacist | 4 | 0.8% |
|  | Technician | 51 | 10.7% |
| Educational level | Elementary degree | 1 | 0.2% |
|  | Intermediate degree | 7 | 1.5% |
|  | Secondary degree | 13 | 2.7% |
|  | Higher degree | 110 | 23.1% |
|  | Bachelor degree | 337 | 70.6% |
|  | Missing | 9 | 1.9% |
| Years of Experience | <5 years | 188 | 39.4% |
|  | 5 – 10 years | 155 | 32.5% |
|  | 10> years | 126 | 26.4% |
|  | Missing | 8 | 1.7% |

Nurses represented a majority (240, 50.3%) with respect to job category. Next were physicians (156, 32.7%) who were followed by Technicians (51, 10.7%); Specialists (15, 3.1%); Non-HW (8, 1.7%) and the least were Pharmacists (4, 0.8%), consecutively, with 3(0.6%) missing records.

The educational level reflected a gradual increase approaching higher learning degrees. The least count was seen in Elementary degree (1, 0.2%); Intermediate degree (7, 1.5%); Secondary degree (13, 2.7%); Higher degree (110, 23.1%) and Bachelor degree were a majority (337, 70.6%), whilst, 9 were missing records (1.7%). Years of work experience were reflected in three ranges <5 years (188, 39.4%); 5 – 10 years (155, 32.5%) and 10> years (126, 26.4%) and 8 missing records (1.7%).

Table 2 showed the information about the viral diseases and their associated signs and symptoms as queries, participants answers, frequencies and percentages. Q1 shows that the majority (444, 93.1%) of participants got the right answer, regarding knowledge of MERS-CoV (Middle east respiratory syndrome).

**Table 2.** Information about the viral diseases (n=477)

| **Table 2. Information about the viral diseases (n=477)** |  |  |  |
| --- | --- | --- | --- |
| **Query** | **Answer** | **Frequency** | **%** |
| **Q 1: What does MERS-CoV stand for?:** |  |  |  |
|  | Middle east respiratory syndrome | 444 | 93.1% |
|  | Moderate encephalitis rapid syndrome | 3 | 0.6% |
|  | Middle east respiratory SARS | 15 | 3.1% |
|  | No idea | 9 | 1.9% |
|  | Missing | 6 | 1.3% |
| **Q 2: Do you know how corona virus is transmitted? You can choose more than one answer:** |  |  |  |
|  | Getting involved with the infected patient | 337 | 70.6% |
|  | Droplets from coughing and sneezing | 405 | 84.9% |
|  | Touching equipment and tools of patients, then touching the mouth, nose, and eyes | 304 | 63.7% |
|  | Can be transmitted via camels | 316 | 66.2% |
| **Q 3: How did you hear about MERS-CoV? You can choose more than one answer:** |  |  |  |
|  | TV/radio/newspaper | 290 | 60.8% |
|  | Physician/ health-care worker | 360 | 75.5% |
|  | Internet/social media | 305 | 63.9% |
|  | Ministry of health pamphlets | 211 | 44.2% |
|  | Seminars | 95 | 19.9% |
|  | Others | 15 | 3.1% |
| **Q 4: One of the measurements to prevent getting infected by MERS-CoV in the waiting area is:** |  |  |  |
|  | Avoid touching any surface in waiting rooms | 138 | 28.9% |
|  | Leaving at least a distance of 2 meters between patients in the waiting area | 53 | 11.1% |
|  | Wearing a protective mask | 256 | 53.7% |
|  | Avoid interacting with patients until they are fully recovered | 24 | 5.0% |
|  | Missing | 6 | 1.3% |
| **Q 5: One of the following might be a symptom of MERS-CoV infection:** |  |  |  |
|  | Constipation | 6 | 1.3% |
|  | Diarrhea | 409 | 85.7% |
|  | Fainting spells | 30 | 6.3% |
|  | Nose bleed | 28 | 5.9% |
|  | Missing | 4 | 0.8% |

In Q2, where the participants could choose more than one chosen answer to ‘How corona virus is transmitted’, their answers were in the following order: ‘Droplets from coughing and sneezing is transmitting the virus’ had the first and highest (405, 84.9%); ‘Getting involved with the infected patient’ was second high (337, 70.6%); next record (316, 66.2%) was ‘Can be transmitted via camels’; whilst, ‘touching patients’ equipment and tools’ was the last and least (304, 66.2%).

In Q3, also the participants could choose more than one to answer ‘How did you hear about MERS-COV?’, and their answers were in the following order: ‘Physician/ health-care worker’ had the first and highest (360, 75.5%); ‘Internet/social media’ was second high (305, 63.9%); next record (290, 60.8%) was ‘TV/radio/newspaper’; next (211, 44.2%) was ‘Ministry of health pamphlets’; whilst, ‘Seminars’ and ‘Others’ were the last and least, with records of (95, 19.9%) and (15, 3.1%), respectively.

Q4 showed that the majority (256, 53.7%) of participants got the answer ‘Wearing a protective mask’, regarding the query ‘one of the measurements to prevent getting infected by MERS-CoV in the waiting area’. Next, were those who chose (138, 28.9%) the answer ‘Avoid touching any surface in waiting rooms’. Whilst, those who chose (53, 11.1%) the answer ‘Leaving at least a distance of 2 meters between patients in the waiting area’, and (24, 5%) the answer ‘Avoid interacting with patients until they are fully recovered’ were the last, and there were (6, 1.3%) missing.

Q5 reflects answers of participants to the query ‘one of the following might be a symptom of MERS-CoV infection’ where a majority (409, 85.7%) of participants gave ‘Diarrhea’ as an answer. Next, (30, 6.3%) were those who gave ‘Fainting spells’ as an answer, whilst those who recorded (28, 5.9%) gave the answer ‘Nose bleed’. Those who gave ‘Constipation’ as an answer were the least (6, 1.3%) and 4 (0.8%) were missing.

Table 3 reflects the use of protective equipment profile. In Q1, choosing ‘The right order to put on personal protective equipment is’, a majority (373, 78.2%) chose ‘Wash hands, put on gown, put on mask, wear eye protection, put on gloves’. The rest chose ‘Wash hands, put on mask, put on gloves, wear eye protection, put on gown’ with a record of 50 (10.5%); ‘Wash hands, put on gloves, wear eye protection, put on gown, put on mask’ with a record of 39 (8.2%); ‘Wash hands, wear eye protection, put on gloves, put on gown, put on mask’ with a record of 11 (2.3%); and 4 (0.8%) were missing.

**Table 3.** MERS-CoV transmission and protection (Protective equipment) (n=477)

|  |  |  |  |
| --- | --- | --- | --- |
| **Query** | **Answer** | **Frequency** | **%** |
| **Q 1: Protective equipment profile:** |  |  |  |
| The right order to put on personal protective equipment is: | Wash hands, put on gown, put on mask, wear eye protection, put on gloves | 373 | 78.2% |
|  | Wash hands, put on gloves, wear eye protection, put on gown, put on mask | 39 | 8.2% |
|  | Wash hands, put on mask, put on gloves, wear eye protection, put on gown | 50 | 10.5% |
|  | Wash hands, wear eye protection, put on gloves, put on gown, put on mask | 11 | 2.3% |
|  | Missing | 4 | 0.8% |
| The correct order to take off personal protective equipment is: | Take off gloves, wash hands, take off gown, take off eye protection | 81 | 17.0% |
|  | Take off gloves, wash hands, take off eye protection, take off gown | 59 | 12.4% |
|  | Take off gloves, take off gown, wash hands, take off eye protection | 101 | 21.2% |
|  | Take off gloves, take off eye protection, take off gown, wash hands | 228 | 47.8% |
|  | Missing | 8 | 1.7% |
| **Q 2: n--95 mask profile? (n=477):** |  |  |  |
| The proper way to wear an n--95 mask is: | Make sure it covers your nose and mouth, and check for air leakage during exhalation | 182 | 38.2% |
|  | Make sure it covers your nose, and check for air leakage during exhalation | 25 | 5.2% |
|  | Make sure it covers your mouth, and check for air leakage during exhalation | 5 | 1.0% |
|  | Make sure it covers your nose, mouth, and chin and check for air leakage during exhalation | 255 | 53.5% |
|  | Missing | 10 | 2.1% |
| The first step to remove an n--95 mask properly is: | Lift off the top elastic | 104 | 21.8% |
|  | Lift away from face while holding the elastic | 150 | 31.4% |
|  | Lift off the bottom elastic over your head | 115 | 24.1% |
|  | Clean hands | 96 | 20.1% |
|  | Missing | 12 | 2.5% |

In choosing ‘The correct order to take off personal protective equipment’, the majority (228, 47.8%) chose ‘Take off gloves, take off eye protection, take off gown, wash hands’. The rest chose ‘Take off gloves, take off gown, wash hands, take off eye protection’ with a record of 101 (21.2%); ‘Take off gloves, wash hands, take off gown, take off eye protection’ with a record of 81 (17.0%); ‘Take off gloves, wash hands, take off eye protection, take off gown’ with a record of 59 (12.4%); and 8 (1.7%) were missing.

In Q2, Choosing ‘The proper way to wear an n-95 mask’, the majority (255, 53.5%) chose ‘Make sure it covers your nose, mouth, and chin and check for air leakage during exhalation’. The rest chose ‘Make sure it covers your nose and mouth, and check for air leakage during exhalation’ with a record of 182 (38.2%); ‘Make sure it covers your nose, and check for air leakage during exhalation’ with a record of 25 (5.2%); ‘Make sure it covers your mouth, and check for air leakage during exhalation’ with a record of 1 (1.0%); and 10 (2.1%) were missing.

Whilst, choosing ‘The first step to remove an n-95 mask properly is’, the majority (150, 31.4%) chose **‘**Lift away from face while holding the elastic**’**. The rest chose ‘Lift off the bottom elastic over your head’ with a record of 115 (24.1%); ‘Lift off the top elastic’ with a record of 104 (21.8%); ‘Clean hands’ with a record of 96 (20.1%); and 12 (2.5%) were missing.

Table 4 showed the participants response to queries about MERS-CoV transmission and protection (Hygienic habits and preventive methods). Q1 reflected the participants response answers to ‘Which of the following steps for hand hygiene is correct?’ with nearly similar cumulative records. The highest (440, 92.2%) record was of the answer ‘Before touching a patient’, followed by the second record (417, 87.4%) of the answer ‘After touching a patient’, the third record (394, 82.6%) of the answer ‘After touching patients’ surroundings’, the fourth record (389, 81.6%) of the answer ‘After body fluid exposure risk’ and the fifth record (388, 81.3%) of the answer ‘Before clean/aseptic procedure’.

**Table 4.**
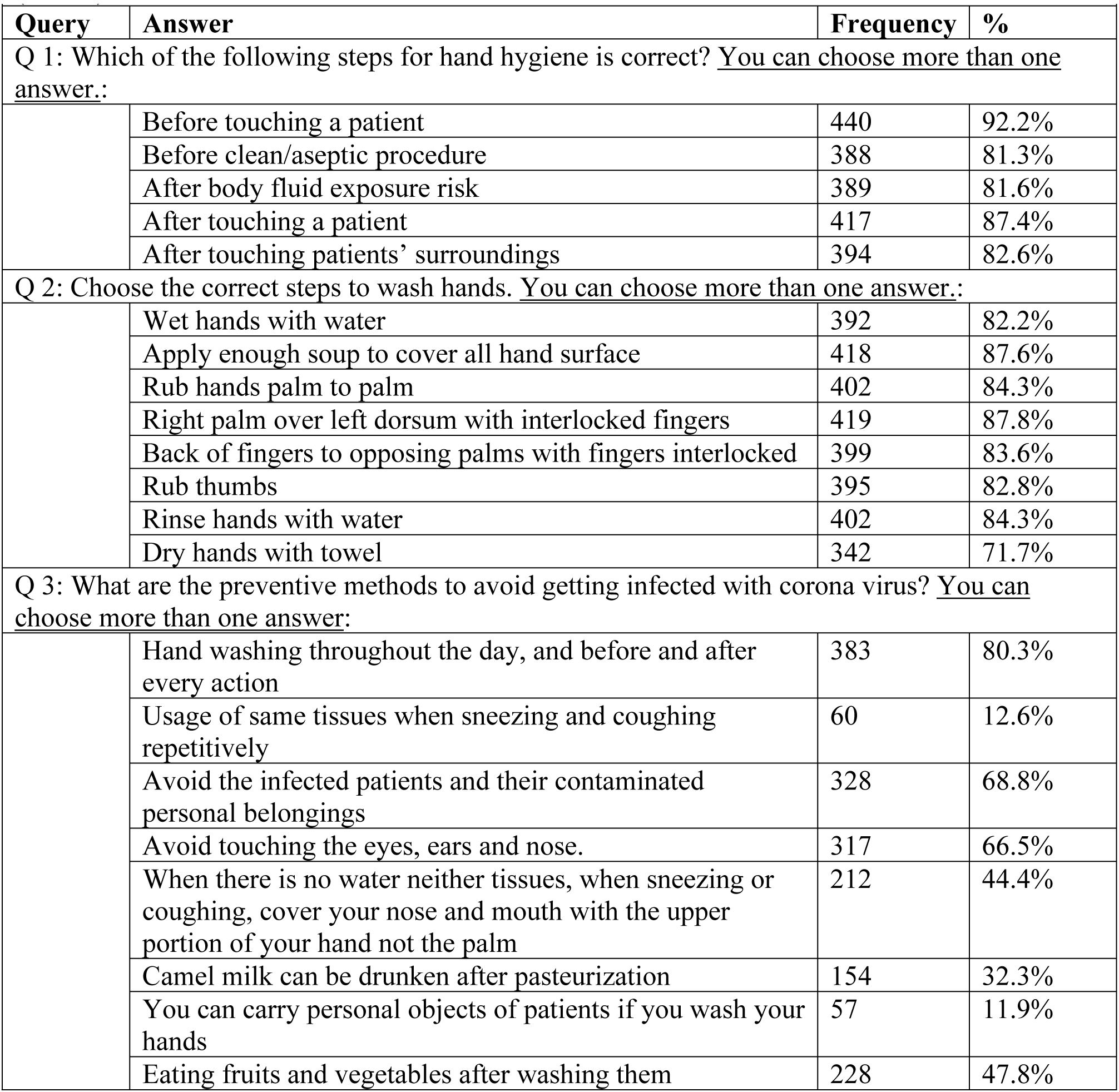
MERS-CoV transmission and protection (Hygienic habits and preventive methods) (n=477)

|  |  |  |  |
| --- | --- | --- | --- |
| **Query** | **Answer** | **Frequency** | **%** |
| Q 1: Which of the following steps for hand hygiene is correct? <u>You can choose more than one answer.:</u> |  |  |  |
|  | Before touching a patient | 440 | 92.2% |
|  | Before clean/aseptic procedure | 388 | 81.3% |
|  | After body fluid exposure risk | 389 | 81.6% |
|  | After touching a patient | 417 | 87.4% |
|  | After touching patients' surroundings | 394 | 82.6% |
| Q 2: Choose the correct steps to wash hands. <u>You can choose more than one answer.:</u> |  |  |  |
|  | Wet hands with water | 392 | 82.2% |
|  | Apply enough soap to cover all hand surface | 418 | 87.6% |
|  | Rub hands palm to palm | 402 | 84.3% |
|  | Right palm over left dorsum with interlocked fingers | 419 | 87.8% |
|  | Back of fingers to opposing palms with fingers interlocked | 399 | 83.6% |
|  | Rub thumbs | 395 | 82.8% |
|  | Rinse hands with water | 402 | 84.3% |
|  | Dry hands with towel | 342 | 71.7% |
| Q 3: What are the preventive methods to avoid getting infected with corona virus? <u>You can choose more than one answer:</u> |  |  |  |
|  | Hand washing throughout the day, and before and after every action | 383 | 80.3% |
|  | Usage of same tissues when sneezing and coughing repetitively | 60 | 12.6% |
|  | Avoid the infected patients and their contaminated personal belongings | 328 | 68.8% |
|  | Avoid touching the eyes, ears and nose. | 317 | 66.5% |
|  | When there is no water neither tissues, when sneezing or coughing, cover your nose and mouth with the upper portion of your hand not the palm | 212 | 44.4% |
|  | Camel milk can be drunken after pasteurization | 154 | 32.3% |
|  | You can carry personal objects of patients if you wash your hands | 57 | 11.9% |
|  | Eating fruits and vegetables after washing them | 228 | 47.8% |

Q2 reflected the participants response answers to ‘Choose the correct steps to wash hands’ with almost similar cumulative records. The first and highest (419, 87.8%) record was of the answer ‘Right palm over left dorsum with interlocked fingers’, followed by the second record (418, 87.6%) of the answer ‘Apply enough soap to cover all hand surface’, the third record (402, 84.3%) of the answer ‘Rinse hands with water’, the fourth record (402, 84.3%) of the answer ‘Rub hands palm to palm’, the fifth record (399, 83.6%) of the answer ‘Back of fingers to opposing palms with fingers interlocked’, the sixth record (395, 82.8%) of the answer ‘Rub thumbs’, the seventh record (392, 82.2%) of the answer ‘Wet hands with water’, and the eighth record (342,71.7%) of the answer ‘Dry hands with towel’.

Q3 showed the chosen answers to ‘What are the preventive methods to avoid getting infected with corona virus?’. The first and highest (383, 80.3%) record was of the answer ‘Hand washing throughout the day, and before and after every action’, followed by the second record (328, 68.8%) of the answer ‘Avoid the infected patients and their contaminated personal belongings’, the third record (317, 66.5%) of the answer ‘Avoid touching the eyes, ears and nose.’, the fourth record (228, 47.8%) of the answer ‘Eating fruits and vegetables after washing them’, the fifth record (212, 44.4%) of the answer ‘When there is no water neither tissues, when sneezing or coughing, cover your nose and mouth with the upper portion of your hand not the palm’, the sixth record (154, 32.3%) of the answer ‘Camel milk can be drunken after pasteurization’, the seventh record (60,12.6%) of the answer ‘Usage of same tissues when sneezing and coughing repetitively’, and the eighth record (57, 11.9%) of the answer ‘You can carry personal objects of patients if you wash your hands’.

Table 5 showed the actions and attitudes towards oneself and towards others during infection of MERS CoV. Q1 reflected the participants response answers to ‘When you suspect that you are being infected with MERS-CoV, what is the one thing you should not do?’ which, showed variable records. The highest (209, 43.8%) record was of the answer ‘If you have fever, shortness of breath, sore throat, cough and sneeze, it is better to stay at home and not to go to work unless 24 hours after the disappearance of these changes.’, followed by the second record (162, 34.0%) of the answer ‘It is tolerable to take over the counter antipyretic and get back to work’, the third record (49, 10.3%) of the answer ‘Avoid any contact with infected patients’ personnel and their stuff of cups, plates or any shared object’, the fourth record (43, 9.0%) of the answer ‘If you were diagnosed with corona infection and no need for hospitalization, follow doctor’s instructions’ and 14 (2.9%) were recorded missing.

**Table 5.** Actions and attitudes towards oneself and towards others during infection of MERS CoV (n=477)

| Query | Answer | Frequency | % |
| --- | --- | --- | --- |
| Q 1: When you suspect that you are being infected with MERS-CoV, what is the one thing you should not do? |  |  |  |
|  | If you have fever, shortness of breath, sore throat, cough and sneeze, it is better to stay at home and not to go to work unless 24 hours after the disappearance of these changes. | 209 | 43.8% |
|  | Avoid any contact with infected patients’ personnel and their stuff of cups, plates or any shared object | 49 | 10.3% |
|  | It is tolerable to take over the counter antipyretic and get back to work. | 162 | 34.0% |
|  | If you were diagnosed with corona infection and no need for hospitalization, follow doctor’s instructions | 43 | 9.0% |
|  | Missing | 14 | 2.9% |
| Q 2: Your actions towards yourself should include: |  |  |  |
|  | Take antipyretics and pain killers | 244 | 51.2% |
|  | Take antibiotics | 152 | 31.9% |
|  | Drink lots of fluid | 400 | 83.9% |
|  | Rest enough | 389 | 81.6% |
|  | Eat healthy food | 362 | 75.9% |
| Q 3: Your action towards others should include: |  |  |  |
|  | Wash your hands routinely | 407 | 85.3% |
|  | Use disposable tissues when coughing and sneezing | 367 | 76.9% |
|  | Avoid hand shaking. | 343 | 71.9% |
|  | Not to share personal equipment like mobile phones | 321 | 67.3% |
|  | Recommend direct contacts to get medications against MERS--COV when you are infected with it. | 221 | 46.3% |

Q2 reflected the participants response answers to ‘When you suspect that you are already infected, your action towards yourself should include’ with variable records. The highest (400, 83.9%) record was of the answer ‘Drink lots of fluids’, followed by the second record (389, 81.6%) of the answer ‘Rest enough’, the third record (362, 75.9%) of the answer ‘Eat healthy food’, the fourth record (244, 51.2%) of the answer ‘Take antipyretics and pain killers’ and the fifth record (152, 31.9%) of the answer ‘Take antibiotics’.

Q3 reflected the participants response answers to ‘Your action towards others should include’, that showed variable records. The highest (407, 85.3%) record was of the answer ‘Wash your hands routinely’, followed by the second record (367, 76.9%) of the answer ‘Use disposable tissues when coughing and sneezing’, the third record (343, 71.9%) of the answer ‘Avoid hand shaking’, the fourth record (321, 67.3%) of the answer ‘Not to share personal equipment like mobile phones’ and the fifth record (221, 46.3%) of the answer ‘Recommend direct contacts to get’.

## Discussion

The levels of awareness concerning MERS CoV has been assessed in staff workers of PSMMC, whose demographic profile reflected similar combinations and responses of the participants, compared with previous studies [23]. Ratios of different groups of HW characteristics like gender, jobs and educational levels reflected a gradual increase approaching higher learning degrees, with Bachelors as a majority, whilst fresh employees were a majority but, the most experienced (>10 years) were the least in number. This situation of demographic group counts order had been noticed before [23], [24].

With respect to information about the viral diseases, the participants’ knowledge regarding basic terminologies like MERS-CoV and viral transmission, the majority got the right answer (Middle east respiratory syndrome). Whilst, the highest score recorded by participants, regarding their multiple choice answer about viral transmission of MERS was the unhygienic spread of coughing and sneezing droplets. Other methods of spread of this virus were also highly appreciated, as has been reported by previous studies [8].

The spreading of information about MERs-COV in society by HW was affirmed by the majority, whilst, spreading of information through internet/social and traditional medias like TV, radio and newspapers were of similar, second option. Ministry of health pamphlets and seminars were of less impact, a situation that demands more attention by the ministry of health in this field.

A majority of the participants stated that wearing a protective mask was one of the measurements to prevent getting infected by MERS-CoV in the waiting area. Whilst, the most effective method to deal with this situation was to avoid the patients until they recover. Other options were also found to be useful measurements with less variable grades. Investigators and other workers in this field found these measures also useful [25].

In evaluating knowledge about associated signs and symptoms of MERS-CoV infection, a majority of the participants chose ‘Diarrhea’ as an answer. On this point, the answer with such an appreciable majority, shouldn’t be overlooked, without some attention. Diarrhea with fever was said to be one of the symptoms of MERS-CoV infection when people were immuno-compromised or organ transplanted, a situation where these patients might not experience the usual route respiratory symptoms [26].

The participants got it quite right in dealing with their protective equipment and showing the right order of putting them on, whilst, they were also correct in taking them off. This reflects that participants were very careful in doing the routine steps before entering their duties in epidemics of seriously infected fields; they were also careful when they completed their jobs and abandoning these protective equipment [22].

Another challenge for these participants assessing their skills abilities in showing the proper way to wear an n-95 mask, specifically used during MERS-CoV infections. In dealing with this type of mask, they were correct in choosing the right steps, which reflects their seriousness and carefulness in entering infected zones, but, they were that quite careful in taking it off, as they put this choice as a second step. Whilst, Attention must be paid to this point, as it might reflect some unjustified hastiness practiced by professionals [20]. Also assessment of the participants reflected their high awareness of infection transmission and protection as precautionary steps for having good hand hygiene experience and practice during rush hours and epidemics. Majorly stating the right timing use reflects the strict adherence to hand hygiene which, is an essential and a basic practice and regime in health fields [20].

In choosing the correct steps in applying and practicing hand wash, the participants were able to get the highest scores which, meant that they understand and are well aware of correct and full hand wash, infection transmission and protection. Appropriate choices and proper practice preferences in medical and hygienic media and fully correct hand wash policy affirms and guarantees healthiness and safety [2], [27].

With respect to additional preventive methods to avoid getting infected with corona virus, they chose five ones with high records. They adhered to the basic rule of hand washing throughout the day, before and after every action whilst, they also preferred to avoid infected patients and their contaminated belongings, their own eyes, ears and nose and to eat fruits only after thoroughly washing them. It was important to mention to use proper compensatory methods to prevent cross infection when sneezing or coughing. Keeping always clean and avoiding sources of infection is always a hygienic routine that guarantees awareness and gratifies with success in health media [2].

Attitudes and actions evaluation of the participants towards themselves and towards others reflected high levels of awareness and knowledge. In certain cases of being sick, generally, people commit serious mistakes and decisions, even within the health field workers. Decision to stay at home with such obvious symptoms of, at least common colds, is a serious one. Also to have your prescription such as antipyretics over the counter and join work is not at all advisable. It is a double hazard and a dangerous behavior, to the patient himself and to his colleagues at the same time. It is an appreciable level of knowledge and good practice that the participants responded correctly to these queries, whilst, few ones might have not understood the rest of queries correctly. With seriously infective viruses such as MERS-CoV, it is mostly crucial to get access to the right treatment and avoid spreading the virus in the vicinity [2].

During serious episodes of community viral infections like MERS-CoV, people should take precautionary steps and build good protective attitudes and actions, towards oneself and towards others in the community. It is quite convincing that participants in the health field have built good and correct awareness to take attitudes of taking lots of fluids, having healthy foods and taking sufficient and beneficial body rest.

Whilst, to help in cutting the vicious circuit of infective body contacts, it is always best attitude towards others to clean hands routinely, use disposable tissues to mask coughing and sneezing, and evade shaking hands. It is highly appreciated that participants in health media should stick to building and adhering to such attitudes [27].

## Conclusions

The majority of the hospital staff was expatriates and the nursing group was the largest which, emphasizes the selective need for Saudization of this health sector. Ministry of health informative pamphlets and seminars about MERS CoV were of less impact, according to the participants opinions, a situation that demanded more attention in this field. Professional HW were quite aware of the basic and newly emergent policies and were very careful of actions before and during their duties in epidemics of seriously infected patient media as those of MERS-CoV.

Attention must be paid by health professionals to using special protective aids, as dealing with them inappropriately might have reflected some unjustified hastiness practiced by them. Strict adherence of HW to correct hand hygiene experience affirmed and guaranteed healthiness and safety whilst, ideal protective attitudes towards themselves and others represented the firm plateau of awareness they had attained in corona virus combat.

## Acknowledgements

We would like to deeply thank Dr. Susan Al Maddah, Director of the Scientific Research Center (SRC) for supporting this research. We would like also to thank Mr. Nasreddien M. Abdo Osman for compiling the manuscript.

## Appendix I

### Imam Muhammed Ibn Saud Islamic University College of Medicine

#### Awareness of MERS—Co Virus Among Staff Members of Prince Sultan Military Medical City in Riyadh, Saudi Arabia

Please take a few minutes to fill out this survey on your awareness of MERS–-Co Virus. We welcome your feedback at the end, and your answers will be kept confidential. Thank you for your participation.

For any further assistance, please feel free to contact us at

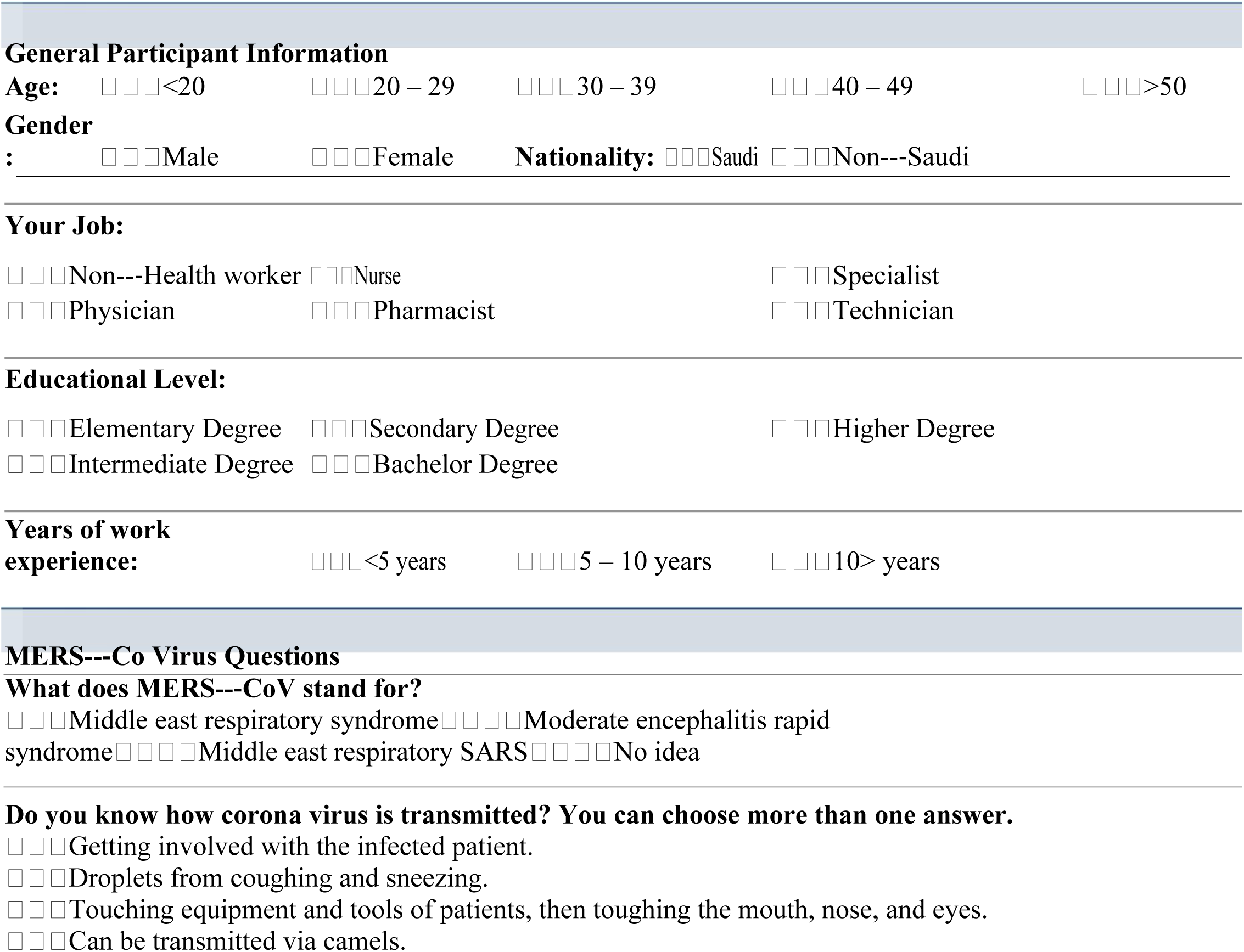
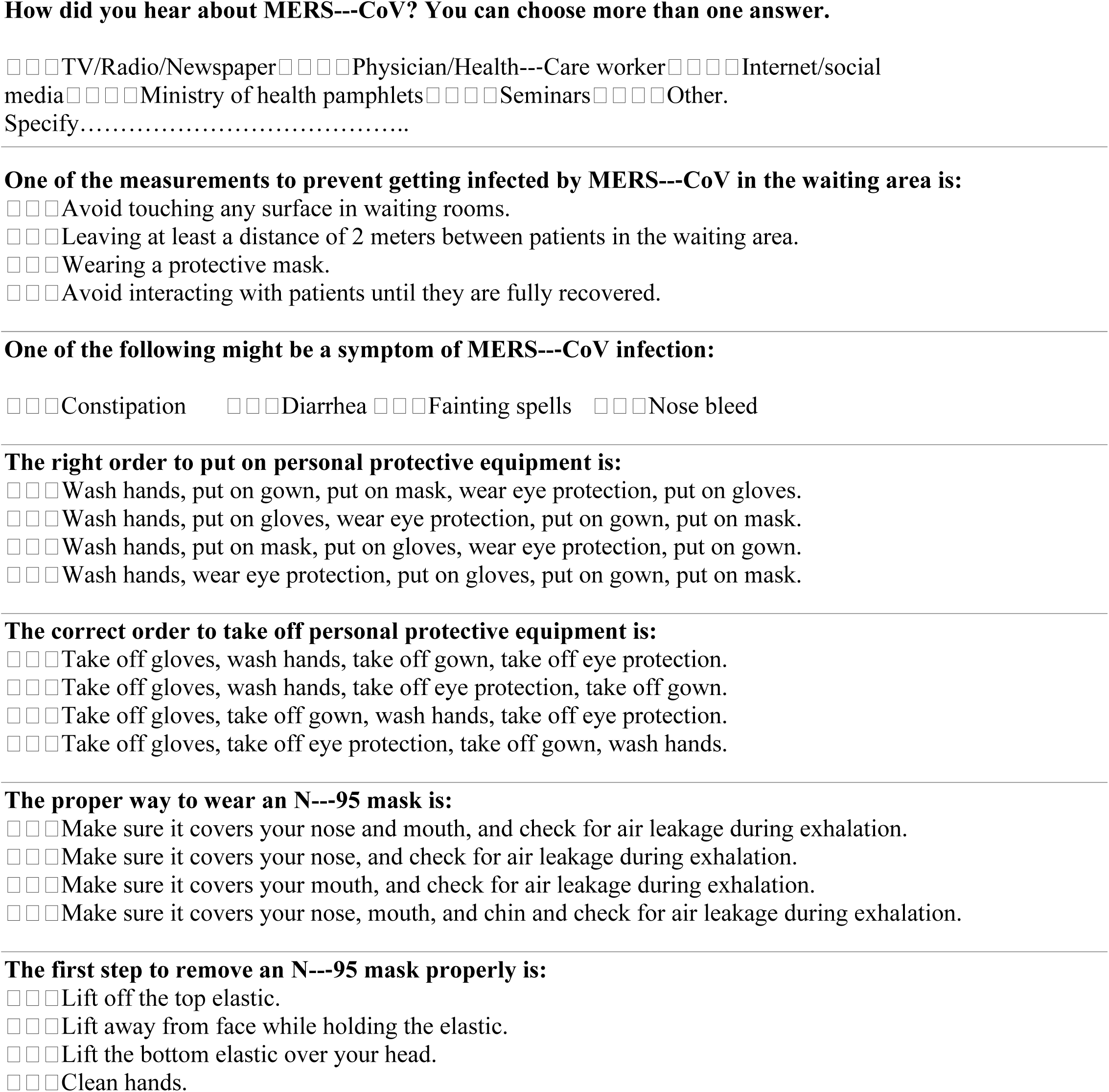
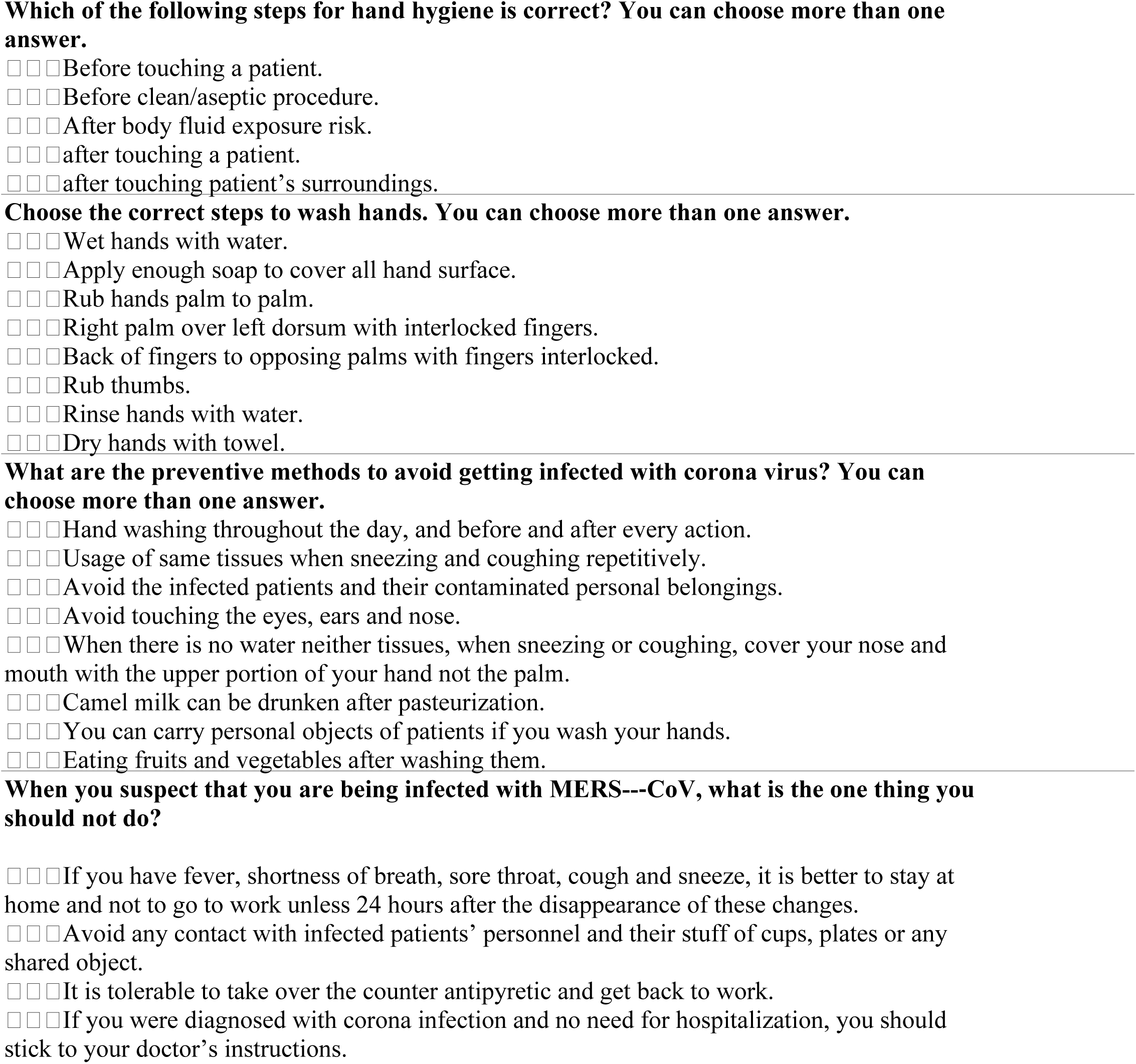
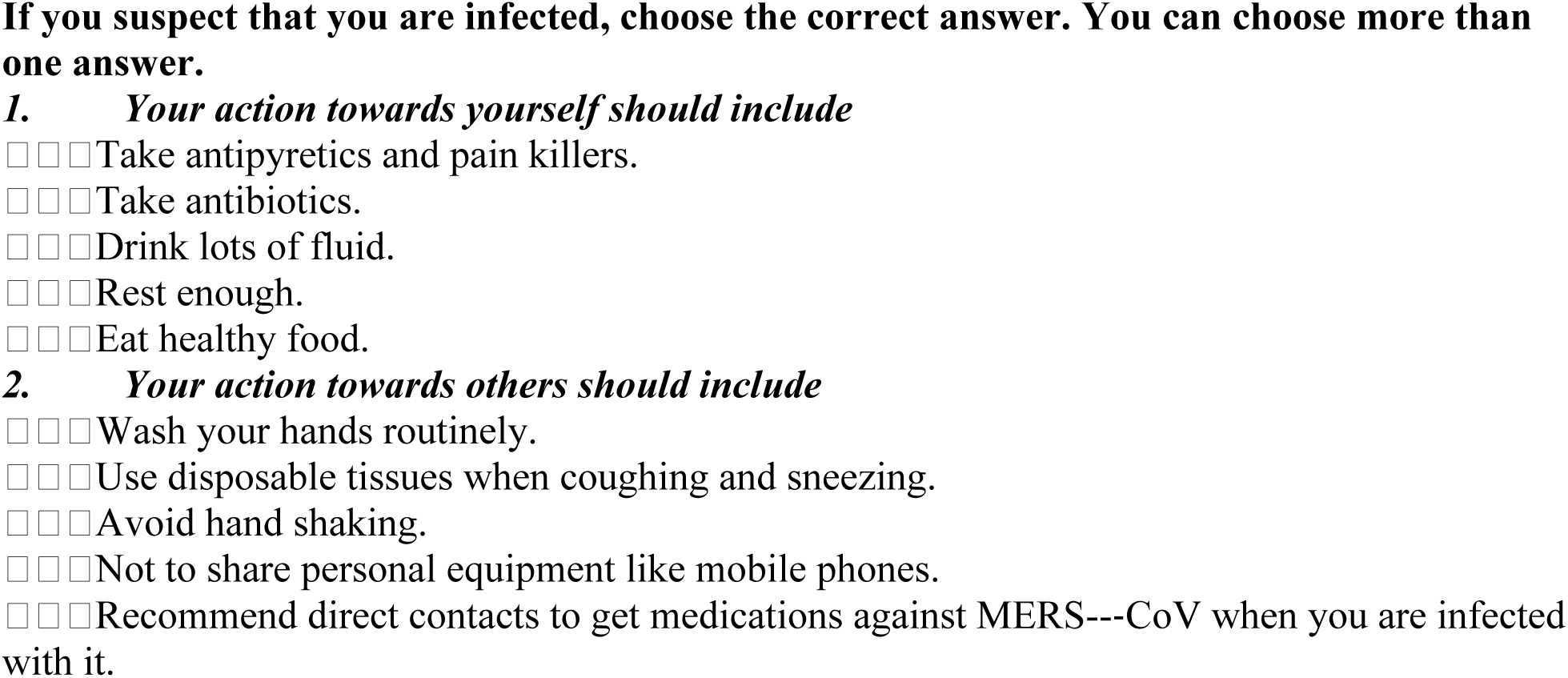

**Thank you very much for your highly appreciated participation in our research**

## References

1. Mohd HA, Al-Tawfiq JA Memish ZA. Middle East Respiratory Syndrome Coronavirus (MERS-CoV) origin and animal reservoir. Virol J. 2016; 13(1):87–93. DOI 10.1186/s12985-016-0544-0

2. World Health Organization (WHO). Middle East respiratory syndrome coronavirus (MERS-CoV)— update. Disease Outbreak News. (http://www.who.int/csr/don/2014_07_14_mers/en/). Updated, 2014.

3. Memish ZA, Mishra N, Olival KJ, Fagbo SF, Kapoor V, Epstein JH, et al. Middle East Respiratory Syndrome Coronavirus in Bats, Saudi Arabia. Emerg Infect Dis. 2013; 19(11): 1819–1823. doi: 10.3201/eid1911.131172. PMCID: PMC3837665

4. Zaki AM, van Boheemen S, Bestebroer TM, Osterhaus AD, Fouchier RA. Isolation of a novel coronavirus from a man with pneumonia in Saudi Arabia. N Engl J Med. 2012;367:1814–20. PMID: 23075143 DOI: 10.1056/NEJMoa1211721 https://www.nejm.org/doi/10.1056/NEJMoa1211721?url_ver=Z39.88-2003&rfr_id=ori:rid:crossref.org&rfr_dat=cr_pub%3dwww.ncbi.nlm.nih.gov

5. WHO. Frequently Asked Questions on Middle East respiratory syndrome coronavirus (MERS-CoV). Updated; 2016. http://www.who.int/csr/disease/coronavirus_infections/faq/en/\

6. Assiri A, McGeer A, Perl TM, Price CS, Al Rabeeah AA, Cummings DA, et al. KSA MERS-CoV Investigation Team. Hospital outbreak of Middle East respiratory syndrome coronavirus. N Engl J Med, 2013(369), pp.407–416. doi: 10.1056/NEJMoa1306742 https://www.ncbi.nlm.nih.gov/pmc/articles/PMC4029105/

7. Drosten C, Seilmaier M, Corman VM, Hartmann W, Scheible G, Sack S, et al. Clinical features and virological analysis of a case of Middle East respiratory syndrome coronavirus infection. Lancet Infect Dis. 2013;13(9): 745–751.

8. World Health Organization (WHO). Middle East respiratory syndrome coronavirus (MERS-CoV). Updated, 2018. http://www.who.int/en/news-room/fact-sheets/detail/middle-east-respiratory-syndrome-coronavirus-(mers-cov)

9. Guery B, Poissy J, el Mansouf L, Séjourné C, Ettahar N, Lemaire X, et al. Clinical features and viral diagnosis of two cases of infection with Middle East Respiratory Syndrome coronavirus: a report of nosocomial transmission. The Lancet 2013; 381(9885): 2265–2272. https://europepmc.org/abstract/med/23727167

10. Center for Diseases Control and Prevention (CDC). MERS-CoV Infection Prevention and Control for Hospitalized Patients. Updated, 2016. http://www.cdc.gov/coronavirus/mers/infection-prevention-control.html

11. Ahmed AE. The predictors of 3- and 30-day mortality in 660 MERS-CoV patients. BMC Infect Dis. 2017; 17: 615–622. DOI: 10.1186/s12879-017-2712-2 https://www.researchgate.net/publication/319642183_The_predictors_of_3-_and_30-day_mortality_in_660_MERS-CoV_patients

12. Rossignol JF. Nitazoxanide, a new drug candidate for the treatment of Middle East respiratory syndrome coronavirus. J Infect Public Health. 2016; 9(3): 227–230. DOI: 10.1016/j.jiph.2016.04.001. https://www.jiph.org/article/S1876-0341(16)30018-1/fulltext

13. Stachulski AV, Santoro MG, Piacentini S, Belardo G, Frazia S, Pidathala C, et al. Second-generation nitazoxanide derivatives: thiazolides are effective inhibitors of the influenza A virus. Future Med Chem. 2018; 10(8): 851–862. https://www.future-science.com/doi/10.4155/fmc-2017-0217

14. Tran K, Cimon K, Severn M, Pessoa-Silva CL, Conly J, (Adapted). Aerosol Generating Procedures and Risk of Transmission of Acute Respiratory Infections to Healthcare Workers: A Systematic Review. CADTH Technology Overviews. 2013; 3(1): 1–3. https://cadth.ca/sites/default/files/aerosol-generating-procedures-and-risk-of-transmission-of-acute-respiratory-infections-a-systematic-review.pdf

15. Breban R, Riou J, Fontanet A. Interhuman transmissibility of Middle East respiratory syndrome coronavirus: estimation of pandemic risk. The Lancet 2013; 382(9893): 694–699. https://www.thelancet.com/journals/lancet/article/PIIS0140-6736(13)61492-0/ppt

16. Omrani AS, Saad MM, Baig K, Bahloul A, Abdul-Matin M, Alaidaroos AY, et al. Ribavirin and interferon alfa-2a for severe Middle East respiratory syndrome coronavirus infection: a retrospective cohort study. Lancet Infect Dis. 2014; 14(11): 1090–1095. https://www.thelancet.com/journals/laninf/article/PIIS1473-3099%2814%2970920-X/fulltext

17. Madani TA, Althaqafi AO, Alraddadi BM. Infection prevention and control guidelines for patients with Middle East Respiratory Syndrome Coronavirus (MERS-CoV) infection. Saudi Med J. 2014; 35 (8): 879–913. http://citeseerx.ist.psu.edu/viewdoc/download?doi=10.1.1.889.187&rep=rep1&type=pdf

18. Wuerz R, Fernandes CM, Alarcon J. Inconsistency of emergency department triage. Emergency Department Operations Research Working Group. Ann Emerg Med. 1998; 32(4):431–435. DOI: https://doi.org/10.1016/S0196-0644(98)70171-4 https://www.sciencedirect.com/science/article/pii/S0196064498701714

19. Gerdtz M, Bucknall T. Australian triage nurses’ decision-making and scope of practice. Aust J Adv Nurs. 2000 Sep-Nov;18(1):24–33. https://www.ncbi.nlm.nih.gov/pubmed/11878360

20. Centers for Disease Control and Prevention (CDC). CDC announces first case of Middle East Respiratory Syndrome Coronavirus infection (MERS) in the United States. 2014. https://www.cdc.gov/media/releases/2014/p0502-US-MERS.html

21. Command and Control Center (CCC). Infection Prevention and Control Guidelines for the Middle East CCC Respiratory Syndrome Coronavirus (MERS-CoV). Scientific Advisory Board, 4rth. Edition. 2017. Command and Control Center, Ministry of Health, Kingdom of Saudi Arabia. https://www.moh.gov.sa/CCC/StaffRegulations/Corona/Documents/Guidelines%20MERS-CoV.pdf

22. Centers for Disease Control and Prevention (CDC). Recommendations for Infection Prevention and Control for *Candida auris*. 2018. Centers for Disease Control and Prevention. National Center for Emerging and Zoonotic Infectious Diseases (NCEZID). Division of Foodborne, Waterborne, and Environmental Diseases (DFWED). https://www.cdc.gov/fungal/candida-auris/c-auris-infection-control.html

23. Pollard E, Hirsh W, Williams M, Buzzeo J, Marvell R, Tassinari A, et al. Understanding employers’ graduate recruitment and selection practices. BIS Research Paper 231. Department for Business, Innovation and Skills, 2015, London, UK. http://publications.aston.ac.uk/29398/1/Employers_graduate_recruitment_and_selection_practices.pdf

24. Al-abdullah N, Kaki R, Almutairi O, Ajaj H, Alamoudi A, Markuobi M, et al. Assessment of the Awareness of Middle East Respiratory Syndrome-Coronavirus Infection In Saudi Arabia: A Cross-Sectional Survey. Internet J Infect Dis. 2016; 15(1):1–8. DOI: 10.5580/IJPH.46719. http://ispub.com/IJID/15/1/46719

25. Sarmiento Guede JR, de Esteban Curiel J, Antonovica A. Viral communication through social media: analysis of its antecedents. Revista Latina de Comunicación Social. 2017; 72: 69–86. http://www.revistalatinacs.org/072paper/1154/04en.html DOI: 10.4185/RLCS-2017-1154

26. Gompf SG, Jones LN, Davis ChP. Middle East Respiratory Syndrome Coronavirus Infection (MERS-CoV Infection). medicinenet, 2016. Accessed July 2018. https://www.medicinenet.com/mers_middle_east_respiratory_syndrome/article.htm#where_can_people_get_more_information_about_mers-cov

27. MOH (Ministry of Health). Specific Precautions against the Middle East Respiratory Syndrome-Coronavirus (MERS-COV). Ministry of Health – Kingdom of Saudi Arabia, 2018. https://www.moh.gov.sa/en/Hajj/HealthGuidelines/HealthGuidelinesDuringHajj/Pages/MERS-COV.aspx

